# Adenosine refines the balance of spermatogonial proliferation and differentiation

**DOI:** 10.64898/2026.10.04.756557

**Authors:** Pengxiang Wang, Duong Nguyen, Cuiying Xiao, YinJiao Xu, Bingjie Hu, Zhaoran Zhang, Marc Reitman, Yuan Wang

## Abstract

The balance between spermatogonial proliferation and differentiation is essential for sustaining male fertility, yet the contribution of cellular metabolism remains poorly understood. Here, we showed that adenosine levels increased significantly during spermatogonial differentiation and promoted the formation of differentiating spermatogonia. Using a conditional knockout mouse model in which *Ahcy*, a gene encoding an enzyme in adenosine synthesis, is deleted from spermatogonia, we found that disruption of adenosine generation blocked spermatogonial differentiation, resulting in a transient delay in spermatogenesis. We further demonstrated that adenosine exerted its effects partially through ADORA2A- and ADORA2B-mediated signaling. Downregulating these receptor activities by specific inhibitors in spermatogonial culture or by genetic ablation in mice led to increased spermatogonial proliferation. Taken together, our study identified adenosine as a metabolic signal in refining the balance between spermatogonial proliferation and differentiation, linking cellular metabolism to critical developmental timing and stem cell fate decisions.

## Introduction

Spermatogenesis is a postnatal process of germ cell development that continuously generates millions of sperm per day to sustain male fertility. During spermatogenesis, spermatogonial stem cells (SSCs) maintain a pool of undifferentiated spermatogonia, which must transiently amplify before undergoing irreversible differentiation ^1^. Uncontrolled spermatogonial proliferation may lead to tumor formation, while insufficient spermatogonial expansion will compromise sperm production and cause male infertility or subfertility ^1, 2^. Properly balanced spermatogonial proliferation and differentiation require a tightly coordinated network of signaling molecules (*i.e.,* GDNF, FGF, and differentiation cue, retinoic acid/RA) and transcriptional regulators, including stemness marker PLZF (*aka.* ZBTB16) and differentiation marker STRA8 ^2^. Additionally, metabolic changes are increasingly recognized as key players in spermatogonial differentiation and germ cell development. For example, early work by *Nakamura et al*., demonstrated that spermatocytes in rodents cannot utilize glucose efficiently and rather use pyruvate and lactate for ATP production^3, 4^. Recent findings have also highlighted that bioenergetic preference shifts from glycolysis to mitochondrial metabolism during spermatogonial differentiation ^5–8^. It remains unclear whether additional metabolic changes impact germ cell development.

Adenosine is an intermediate metabolite that serves as a building block for nucleic acid synthesis and ATP production. Adenosine may be generated from various metabolic pathways. In cells, adenosine may be produced from adenine nucleotide (*i.e.*, AMP) by two 5’-nucleotidases, NT5C1a and NT5C2 ^9, 10^. In addition, adenosine can also be generated by S-adenosylhomocysteine hydrolase (AHCY) from S-adenosylhomocysteine (SAH) in the methionine cycle ^9, 10^. The level of adenosine may be further regulated by other enzymes in catabolic pathways. For example, adenosine kinase may phosphorylate adenosine, converting it to AMP and reducing cellular adenosine levels ^9, 10^.

Although the cellular adenosine concentration is relatively low, it fluctuates in response to physiological stimuli and pathological stresses. In these cases, adenosine acts as a signal molecule by activating four distinct adenosine receptors, *i.e*., ADORA1, ADORA2A, ADORA2B, and ADORA3 ^11, 12^. These receptors are widely expressed and have been implicated in several biological functions ^9, 11^. For example, adenosine may be incorporated into cAMP and regulate PKA (*aka.* Protein Kinase A) activities by altering cAMP levels ^13^. Alternatively, adenosine can increase cAMP levels through ADORA2A in NK cells ^14^. Adenosine can further activate MEK-ERK (*aka.* MARK/ERK kinase-extracellular signal-regulated kinase) signaling pathways to regulate cell proliferation and neural stem cell differentiation *via* ADORA1 ^15^.

In this study, using targeted metabolomics, we found significantly elevated adenosine levels in differentiating spermatogonia. Altering adenosine levels or inhibiting adenosine receptor activity affected the balance between spermatogonial proliferation and differentiation, both in spermatogonial culture and during *in vivo* germ cell development. Our work thus provides direct evidence that adenosine-mediated pathways play instructive roles in spermatogonial proliferation and differentiation, establishing cellular metabolism as an active player in regulating critical developmental transitions in spermatogenesis.

## Results

### Adenosine promotes spermatogonial differentiation

We previously revealed that the active pentose phosphate pathway is crucial for spermatogonial differentiation ^5^. The pentose phosphate pathway generates ribose-5’-phosphate for nucleotide and nucleic acid synthesis. In an effort to identify the metabolic changes during spermatogonial differentiation, we conducted targeted metabolic profiling on undifferentiated and differentiating spermatogonia induced by RA for 48 hours (Supplemental Fig. S1A-F). We examined the changes of about 200 metabolites, including those involved in nucleic acid and amino acid metabolism. In total, 108 metabolites were detected, and 12 metabolites showed differential levels between undifferentiated and differentiating spermatogonia (Fig. 1A, Supplemental Table S1, and Supplemental Fig. S1F). Among them, the adenosine level was significantly increased in differentiating spermatogonia (Fig. 1A and Supplemental Table S1). We thus assessed the impact of adenosine on spermatogonial differentiation. We found that adenosine treatment during RA-induced differentiation increased the percentage of KIT+ differentiating spermatogonia (Fig. 1B). Consistently, transcript levels of differentiation markers, *Stra8* and *Kit,* were upregulated in the RA-induced group with adenosine addition (Fig. 1C).

**Fig. 1.**
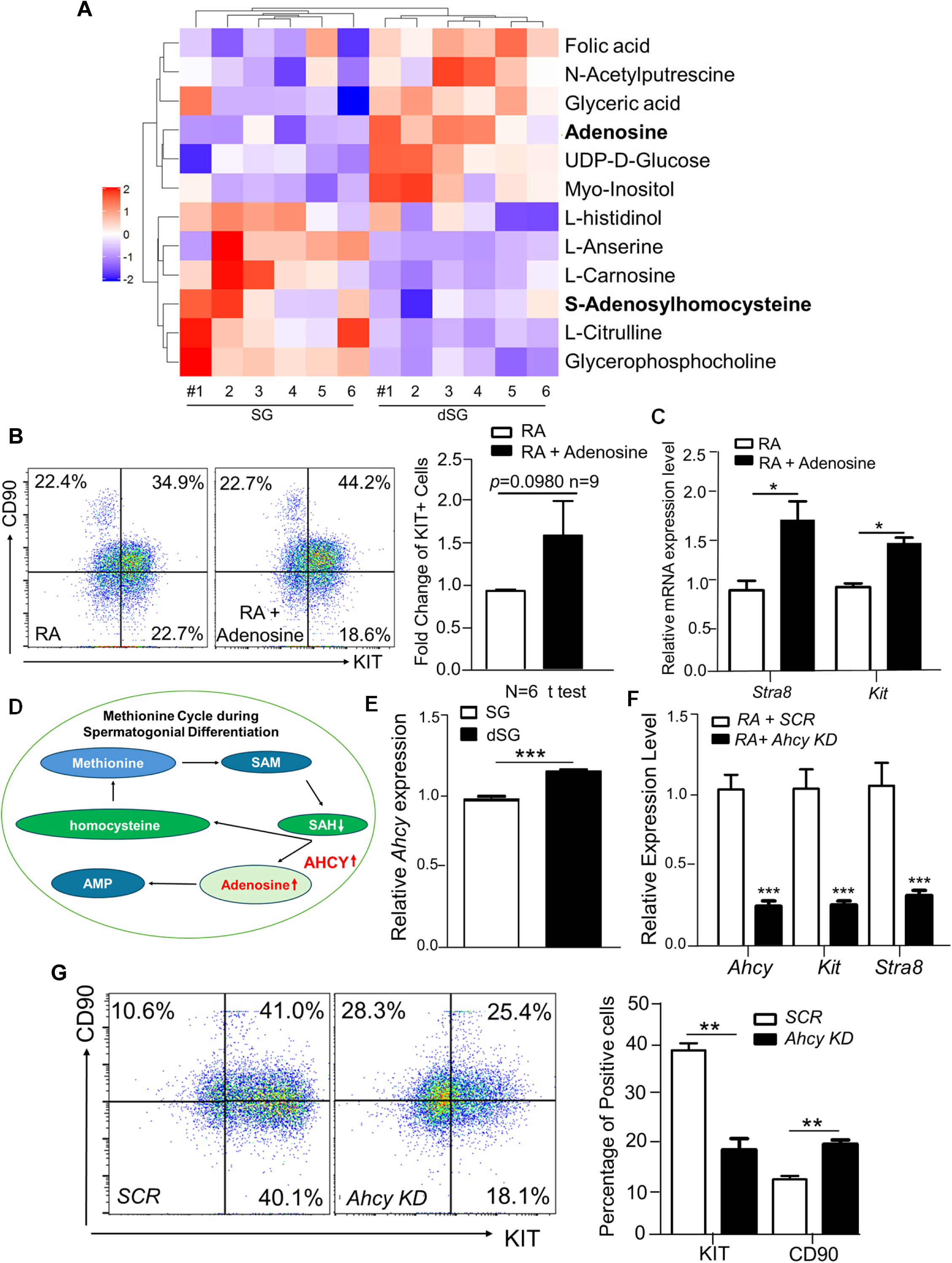
Adenosine promotes spermatogonial differentiation. (**A**) Heatmap of metabolites with differential levels in cultured spermatogonia (SG) and differentiating spermatogonia induced by RA (dSG). Six biological replicates per group. The scale shows the fold change in metabolite levels between dSG and SG. (**B**) Flow cytometry analyses of spermatogonia induced with RA with or without adenosine treatment. A bar graph on the right shows the fold changes in percentage of KIT+ differentiating spermatogonia upon adenosine treatment. Data presented as mean ± SEM, calculated from 9 biological replicates. (**C**) Transcript levels of differentiation markers were analyzed by real-time RT-PCR in RA-induced differentiating spermatogonia. (**D**) Diagram of the methionine cycle, in which SAH is catalyzed into adenosine by AHCY. Please note that SAH decreased, whereas adenosine and AHCY increased, during the differentiation of spermatogonia induced by RA. (**E**) *Ahcy* transcript levels in undifferentiated and differentiating spermatogonia were analyzed by real-time RT-PCR analyses. (**F**) Real-time RT-PCR analyses on RA-induced differentiating spermatogonia with scrambled shRNA (*SCR*) or shRNA against *Ahcy (Ahcy KD)*. (**G**) The percentages of RA-induced KIT+ differentiating spermatogonia and CD90+/KIT-undifferentiated spermatogonia were analyzed by flow cytometry. Left panel: representative flow cytometry graphs. Right panel: bar graph shows the averaged percentage of KIT+ differentiated spermatogonia. (**C, E, F, G**) Data were shown as mean ± SEM. N=3. *: *p* < 0.05; **: *p* <0.01; ***: *p* < 0.001.

From our metabolic analyses, we found that S-adenosylhomocysteine (SAH) was also downregulated (Fig. 1A and Supplemental Table S1). In cells, SAH can be catalyzed by AHCY into adenosine and homocysteine (Fig. 1D)^16^. Interestingly, *Ahcy* transcript levels were upregulated during spermatogonial differentiation (Fig. 1E). We further assessed whether AHCY is required for spermatogonial differentiation by knocking it down using RNA interference in cultured spermatogonia. Indeed, AHCY downregulation decreased *Stra8* and *Kit* expression. KIT+ differentiating spermatogonia induced by RA were downregulated accordingly, while CD90+/KIT-undifferentiated population was increased in the *Ahcy* knockdown group (Fig. 1F & G; Supplemental Fig. S1G & H). Taken together, these data suggest that downregulating adenosine synthesis blocks spermatogonial differentiation.

### Deficient adenosine synthesis leads to delayed spermatogenesis

We next sought to assess the physiological requirement of adenosine during spermatogenesis. An *Ahcy* conditional knockout allele was generated by flanking exon 2 with two *LoxP* sites (Supplemental Fig. S2A-B). We utilized a germ cell-specific *Stra8-Cre* driver ^17^ to delete *Ahcy* from spermatogonia. Homozygote *Ahcy* mutant mice (*i.e., Ahcy^f/f^; Cre*) were obtained by crossing *Ahcy^f/f^* female with *Ahcy^f/+^;Stra8-Cre* male mice. We confirmed the conditional deletion of AHCY by CRE recombinase from DDX4+ germ cells in *Ahcy^f/f^; Cre* mice by immunohistofluorescence (IHF) assays (Fig. 2A).

**Fig. 2.**
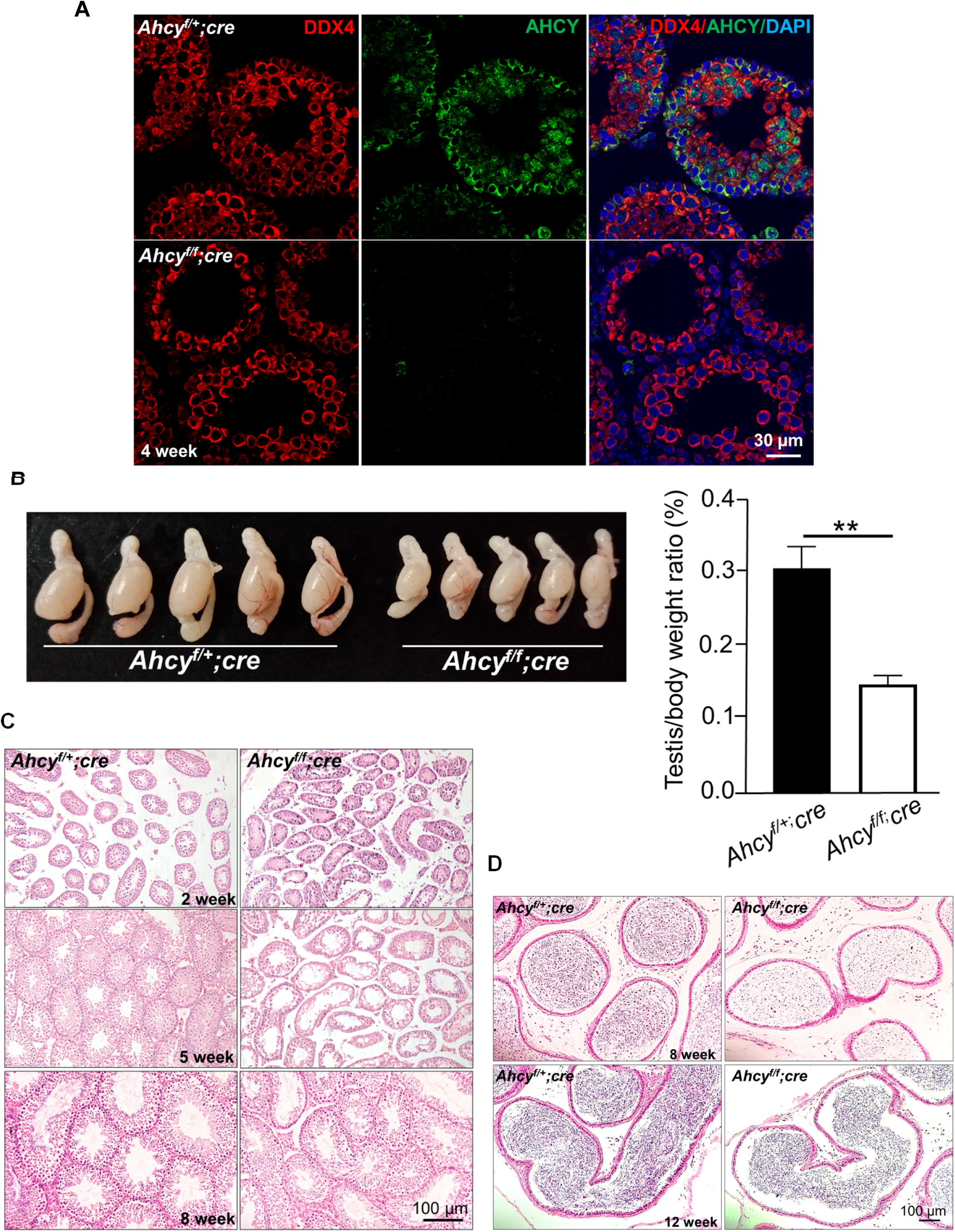
Conditional *Ahcy* deletion from spermatogonia leads to delayed spermatogenesis. (**A**) IHF was performed on testes from 4-week-old mice using antibodies against STRA8 and AHCY, followed by DAPI counterstaining. AHCY signals were not detectable in germ cells upon its conditional deletion from spermatogonia. (**B**) The morphology of testes and epididymis from *Ahcy^f/f^; Cre* mice and wild-type littermate controls at 4 weeks. A bar graph on the right shows the testes-to-body-weight ratio for these mice. Data were shown as mean ± SEM. **: *p* <0.01. N=5. (**C**) Histological studies on testis sections from *Ahcy^f/f^; Cre* mice and littermate controls at postnatal 2, 5, and 8 weeks. (**D**) Histological studies on the epididymis from *Ahcy^f/f^; Cre* mice and littermate controls at postnatal 8 and 12 weeks.

In mice, prospermatogonia develop during the embryonic stage and then become undifferentiated spermatogonia after birth ^2^. The first wave of spermatogenesis occurs during the first 6 weeks of postnatal development. A small fraction of undifferentiated spermatogonia develop into SSCs to support steady-state spermatogenesis in adulthood ^2^. Spermatogonia and SSCs are located in the basal compartment, while spermatocytes and spermatids move toward the center (*i.e.,* adluminal compartment) of seminiferous tubules. Spermatocytes from the first wave of spermatogenesis can be detected before postnatal day (P) 14, while haploid spermatids form around P20 ^2^.

Compared to their littermate controls, the *Ahcy^f/f^; Cre* male mice were fertile. However, the testes appeared to be smaller (Fig. 2B & Supplemental Fig. S2C), suggesting that *Ahcy* deficiency compromised spermatogenesis. We examined germ cell development in these mice using histology at 2 weeks, 5 weeks, and adulthood. We found that the germ cell numbers were drastically reduced at 5-week-old *Ahcy^f/f^; Cre* mice, compared to their littermate controls (Fig. 2C). At 8 weeks, germ cell development was largely recovered in testis, but significantly reduced sperm were detected in epididymis (Fig. 2D & Supplemental Fig. S2D). By 12 weeks, the sperm content was largely back to normal and comparable between *Ahcy^f/f^; Cre* mice and littermate controls (Fig. 2D), albeit that the testes *Ahcy^f/f^; Cre* animals remained smaller (Supplemental Fig. S2C).

We further assessed germ cell composition by IHF with antibodies against SCYP3 (spermatocytes), PRM1 (haploid spermatids), and DDX4 (from spermatogonia to spermatids). In 2-week-old control mice, DDX4+/SYCP3+ spermatocytes were readily detected in the middle of the lumen of seminiferous tubules (Fig. 3A). By contrast, in *Ahcy^f/f^; Cre* mice, DDX4 signals were largely observed at the basal compartment where spermatogonia resided (Fig. 3A). More than 20% of seminiferous tubule sections lacked SYCP3 signal (Fig. 3A and Supplemental Fig. S3A). Delayed spermatogenesis was also observed in mutant mice at 5 weeks old. Seminiferous tubules containing SYCP3+ spermatocytes were reduced in *Ahcy^f/f^; Cre* testis compared to their littermate controls (Fig. 3A and Supplemental Fig. S3B). In addition, PRM1 staining showed that fewer spermatids were formed at 4 weeks in *Ahcy^f/f^; Cre* mice than in littermate controls (Fig. 3A-B). Similarly, reduced PRM1+ sperm were detected by IHF in epididymis from *Ahcy^f/f^; Cre* mice at 8 weeks (Fig. 3C).

**Fig. 3.**
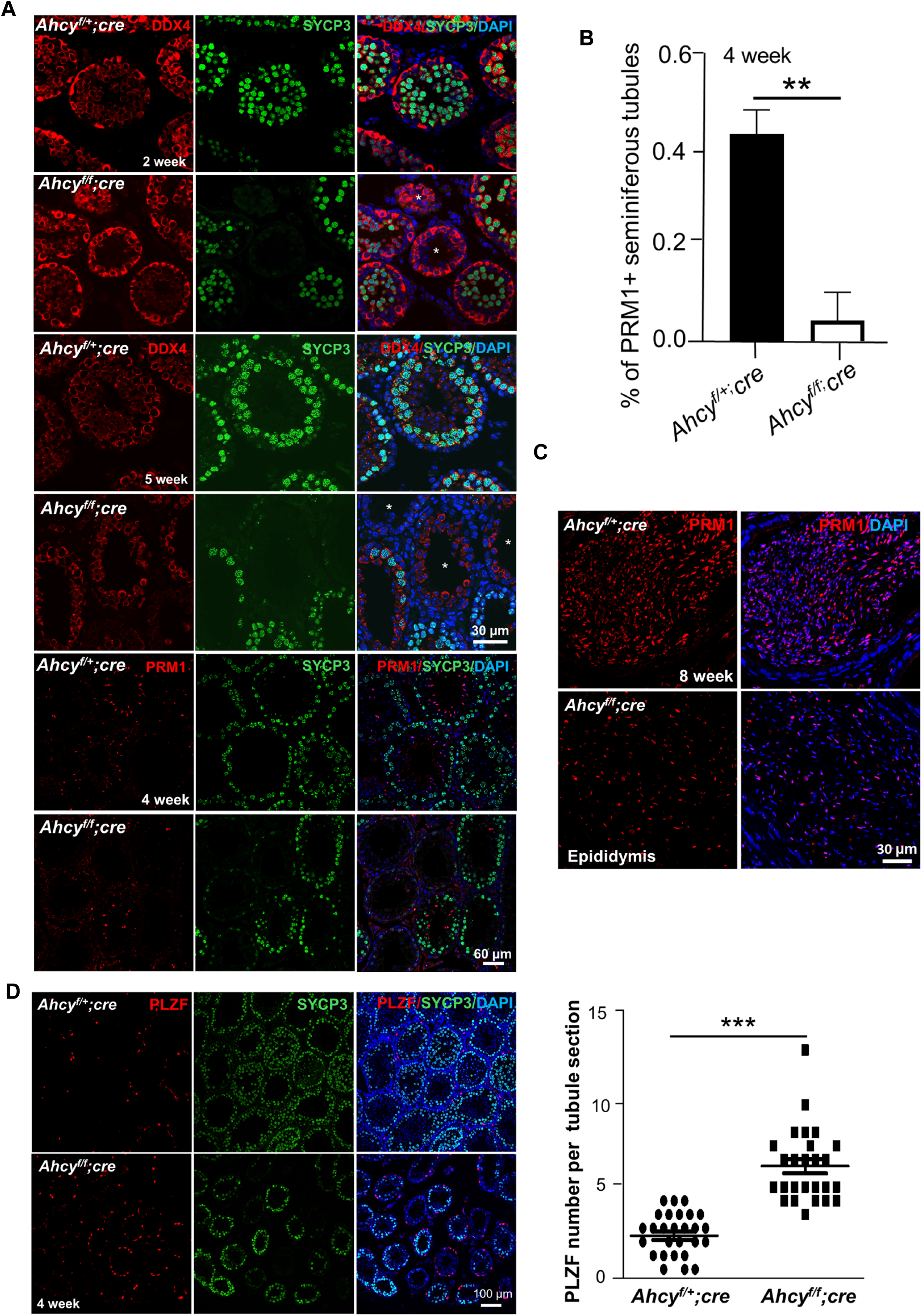
*Ahcy* deficiency reduces spermatogonial differentiation. (**A**) IHF was performed on testes from 2-, 4-, or 5-week-old mice using antibodies against DDX4 and SYCP3, or PRM1 and SYCP3 antibodies, and sections were counterstained with DAPI. White * indicates the seminiferous tubule section without SYCP3+ spermatocytes. (**B**) The bar graph shows the average percentage of seminiferous tubule sections containing PRM1+ cells in testes from 4-week-old mice, calculated from > 8 testis sections. (**C**) IHF was performed on epididymal sections using a PRM1 antibody, and counterstained with DAPI. (**D**) IHF was performed on testes from 4-week-old mice with antibodies against PLZF and SYCP3. Right panel: a bar graph shows the mean number of PLZF+ undifferentiated spermatogonia per seminiferous tubule section. Left panel: Representative images of IHF were shown. (**B, D**). Data were shown as mean ± SEM. **: *p* < 0.01; ***: *p* < 0.001.

Because AHCY downregulation reduced the formation of KIT+ differentiating population induced by RA in cultured spermatogonia (Fig. 1), we examined whether spermatogonial development was altered in *Ahcy^f/f^; Cre* mutant mice at 4 weeks by IHF. We found that PLZF+ undifferentiated spermatogonia increased upon deletion of *Ahcy* from germ cells (Fig. 3D). This finding was further confirmed by flow cytometry analyses. We found that the percentage of CD90+/KIT-undifferentiated spermatogonia in *Ahcy^f/f^; Cre* mutant testes at 4 weeks was increased, while the CD90-/KIT+ differentiating population was decreased (Supplemental Fig. S3C). Taken together, our data suggest that spermatogenesis was transiently delayed due to blocked spermatogonial differentiation and spermatocyte formation upon *Ahcy* deficiency.

### Adenosine negatively regulates spermatogonial proliferation

Adenosine levels were relatively low in undifferentiated spermatogonia but increased during differentiation. This prompted us to examine the effect of additional adenosine on spermatogonial proliferation. Interestingly, we observed that the size of spermatogonial colonies gradually decreased upon the addition of adenosine during a 21-day culture (Fig. 4A-B). Adenosine is a signaling molecule that regulates various biological functions through its 4 receptors, ADORA1, ADORA2a, ADORA2b, and ADORA3 ^11, 12^. To identify which receptor is responsible for the effect of adenosine on spermatogonial development, we examine the expression of these receptors during RA-induced spermatogonial differentiation. We found that receptors ADORA2a and ADORA2b exhibited increased levels in differentiating spermatogonia (Fig. 4C). Levels of these receptors were further potentiated by additional adenosine treatment (Supplemental Fig. S4A), suggesting that ADORA2a and ADORA2b are the receptors that mediate the adenosine effect in spermatogonia. We thus treated undifferentiated spermatogonia with two well-established inhibitors, ZM241385 (an ADORA2a inhibitor) ^18, 19^ and GS6201 (an ADORA2b inhibitor, also known as CVT-6883) ^20^. We found that inhibition of adenosine receptor activity by these inhibitors led to enlarged spermatogonial colonies over time (Fig. 4A-B), suggesting that adenosine negatively regulates spermatogonial proliferation through its receptors.

**Fig. 4.**
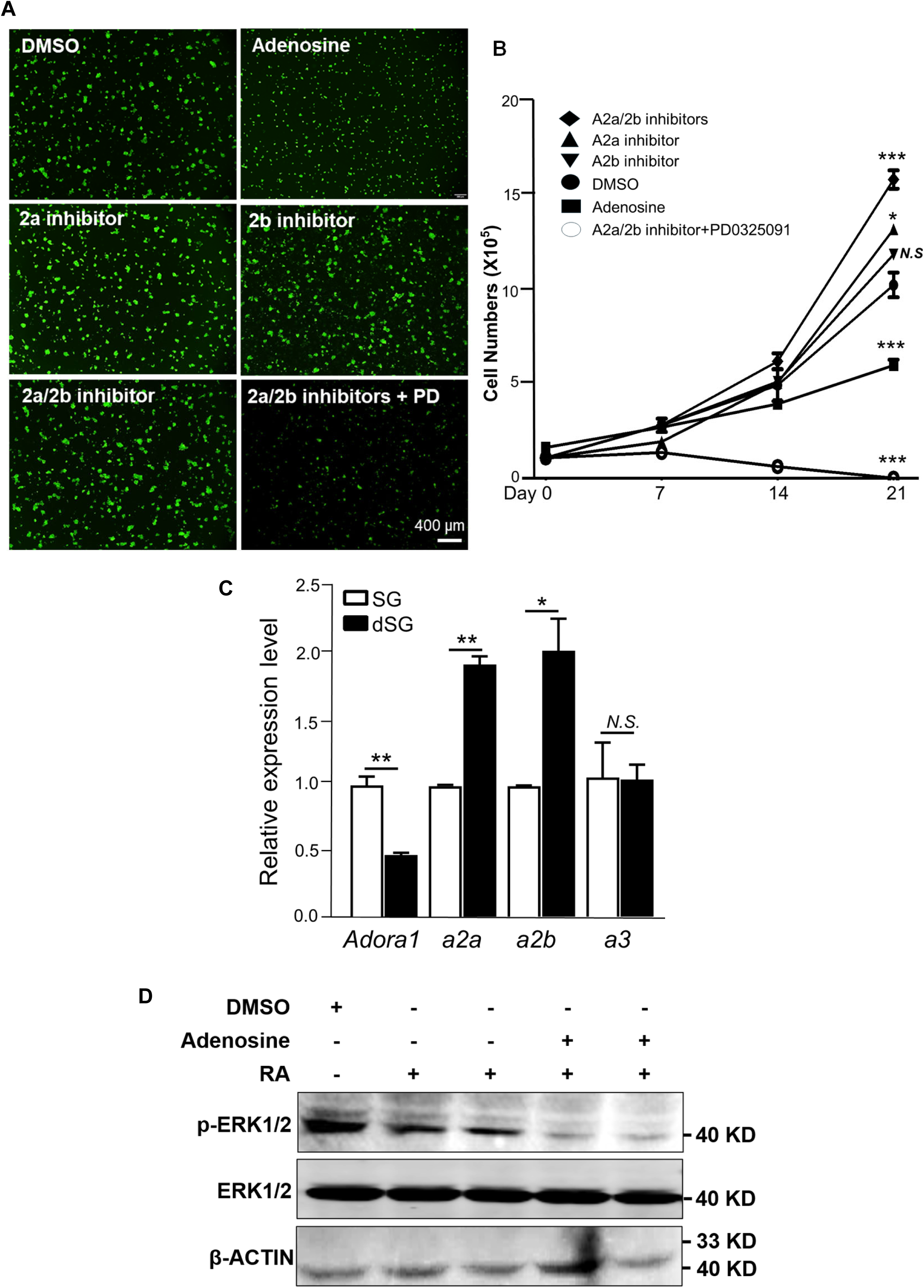
Adenosine inhibition promotes spermatogonia proliferation. (**A**) GFP-expressing spermatogonia were treated with adenosine (100 nM), adenosine receptor inhibitors (2 µM), and with or without an ERK inhibitor (PD, 1 µM). Spermatogonial colonies were visualized under a fluorescent microscope. (**B**) The number of spermatogonia under various treatments was recorded over a 21-day culture period, compared with DMSO controls. (**C**) The expression levels of adenosine receptors in cultured spermatogonia (SG) and differentiating spermatogonia induced by RA (dSG). were analyzed by real-time RT-PCR. (**D**) Western Blotting on spermatogonia treated with various chemicals. The antibodies used are listed on the left. Raw blot images were included as Supplemental Fig. S4B. (**B, C**) Data were shown as mean ± SEM. N = 3. *: *p* < 0.05; **: *p* < 0.01; ***: *p* < 0.001. N.S.: No significance.

The MEK-ERK pathway plays a critical role in spermatogonial proliferation and male germ cell development ^21, 22^. Adenosine is known to modulate MEK-ERK activities through its receptors in Chinese hamster ovary cells and neural stem cells ^15, 23^. Consistent with previous findings ^21, 22^, we found that phosphorylated (p)-ERK1/2 was highly expressed in undifferentiated spermatogonia, downregulated upon RA-induced differentiation, and further reduced with adenosine treatment during this process (Fig. 4D). To investigate whether MEK-ERK is a downstream effector of adenosine in spermatogonia, we treated spermatogonia with both the adenosine inhibitors and PD0325091, an MEK inhibitor. We found that the addition of PD0325091 abolished the promoting effect of adenosine receptor inhibitors (Fig. 4A-B), indicating that MEK-ERK indeed works downstream of adenosine to promote spermatogonial proliferation.

### Deficiency of adenosine receptor activities leads to transiently increased undifferentiated spermatogonia

Adenosine functions as a signaling molecule and may partially regulate biological processes and development *via* its receptors. We found that blocking adenosine receptor activity increased spermatogonial proliferation (Fig. 4). To confirm this finding *in vivo*, we assessed postnatal germ cell development in a mouse model with quad-knockout (QKO) of all four adenosine receptors ^12^. These mice have relatively normal growth, metabolism, and body temperature ^12^. Adult male mice are fertile as well. We analyzed these mice using histology and IHF at P25 and in adulthood. Interestingly, we found that, as in *Ahcy*-deficient mice, QKO mice exhibited transiently delayed spermatogenesis. On postnatal day 25, well-progressed meiosis was observed in wild-type mice with readily detectable SYCP3+ spermatocytes in most seminiferous tubules. By contrast, the QKO testes contained fewer spermatocytes (Fig. 5A-B). Germ cell development was largely restored at 6.5 weeks, with comparable germ cell composition in the testes (Fig. 5A) and sperm formation in the epididymis (Fig. 5C). We further analyzed PLZF+ undifferentiated spermatogonia by IHF at P25. We found that, compared with wild-type controls, QKO mice displayed elevated PLZF+ undifferentiated spermatogonia (Fig. 5D), likely due to increased spermatogonial proliferation and blocked spermatogonial differentiation following deletion of adenosine receptors.

**Fig. 5.**
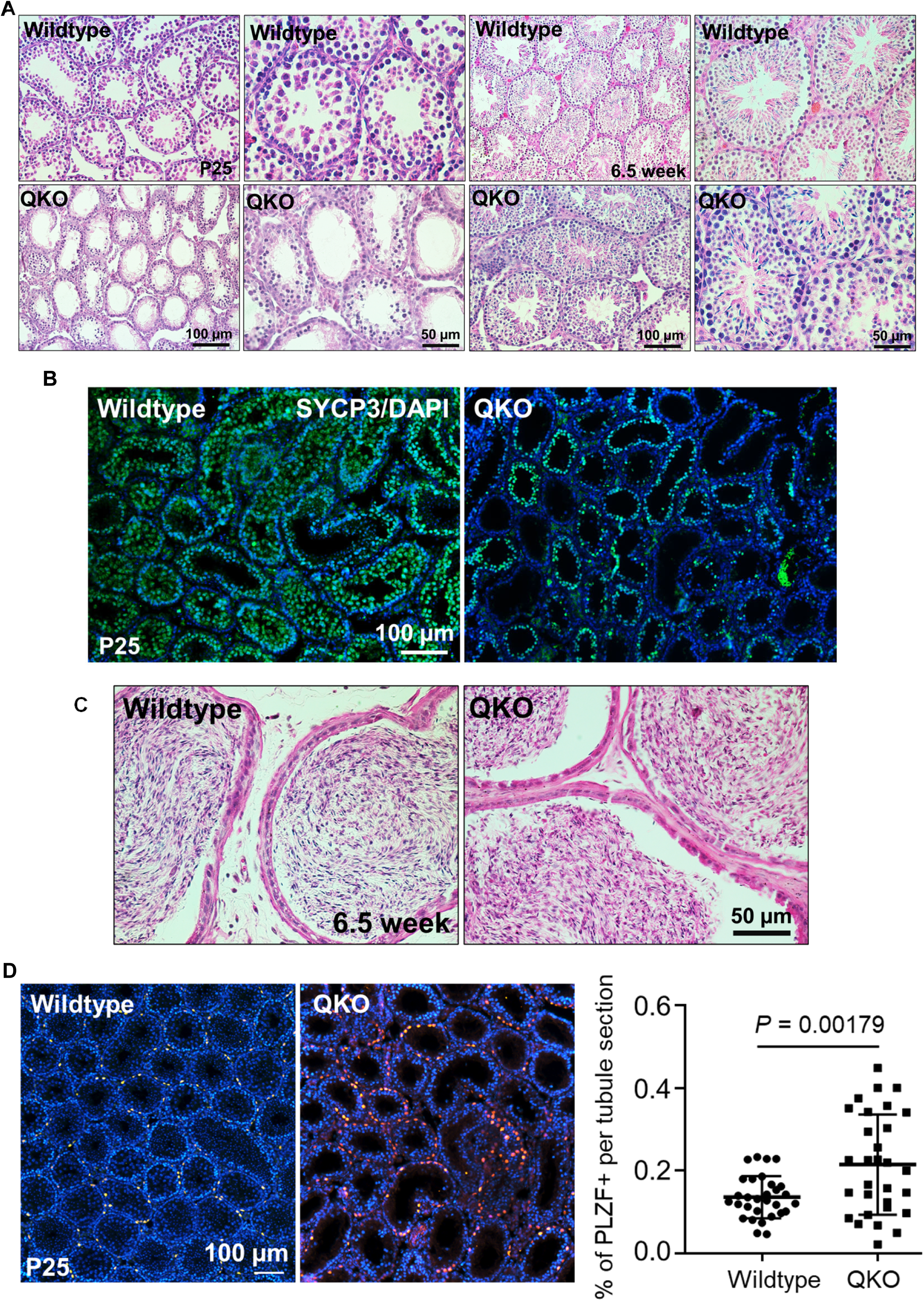
Deletion of adenosine receptors leads to delayed spermatogenesis. (**A**) Histological studies on testis sections from C57BL/6J controls and mice with quad-knockout (QKO) of 4 adenosine receptors at P25 and 6.5 weeks. (**B**) IHF was performed on testes from C57BL/6J control and OKO mice using antibodies against SYCP3 (green), counterstained with DAPI (blue). (**C**) Histological studies on epididymis sections from C57BL/6 control and OKO mice at 6.5 weeks. (**D**) IHF was performed on testes from C57BL/6J control and OKO mice at P25 with antibodies against PLZF (red), counterstained with DAPI (blue). Right panel: a bar graph shows the average PLZF+ undifferentiated spermatogonia per seminiferous tubule section. Left panel: Representative images of IHF were shown. Data were shown as mean ± SEM.

## Discussion

Spermatogonial differentiation is not just a transcriptional or signaling switch. It is tightly coupled to metabolic reprogramming that enables exit from spermatogonial proliferation, entry into meiosis, and germ cell survival. Using targeted metabolomics, we identified adenosine as a regulator for spermatogonial development. Further analyses demonstrated that exogenous adenosine promoted spermatogonial differentiation, whereas inhibition of adenosine synthesis or repression of adenosine receptor activity increased spermatogonial proliferation, blocked spermatogonial differentiation, and delayed spermatogenesis. Our findings thus demonstrate that adenosine functions not merely as a metabolic product, but as an active player that coordinates metabolic reprogramming with cell fate decisions in postnatal male germ cell development.

How does adenosine regulate spermatogonial proliferation and differentiation? First, adenosine is a purine nucleoside that can be converted into ATP to provide energy or into nucleotides as building blocks for cell proliferation. It is thus possible that reduced adenosine levels in *Ahcy* conditional knockout mice affect germ cell health and, consequently, delay spermatogenesis. Notably, exogenous adenosine inhibited undifferentiated spermatogonia,

suggesting a specific requirement for adenosine during differentiation. Second, adenosine is an important regulator of the methionine cycle ^16^, which provides a methyl group for histone, DNA, and RNA modification, thereby affecting the epigenetic status of germ cells ^17, 24, 25^. Indeed, we uncovered that the SAH level was lower in differentiating spermatogonia. Consistently, AHCY, the enzyme that catalyzes SAH to adenosine, was upregulated during spermatogonial differentiation. We further found that conditional deletion of *Ahcy* in spermatogonia resulted in reduced spermatogonial differentiation and delayed spermatogenesis. As SAH generally inhibits S-adenosylhomocysteine (SAM)-dependent methyltransferases ^26^, the conversion of SAH into adenosine decreases SAH level and thus favors DNA and RNA methylation, both of which are increased during spermatogonial differentiation ^17, 24^.

Third, adenosine is a signaling molecule, and it may exert its functions through adenosine receptors ^11^. We observed that the adenosine receptors A2a and A2b were upregulated during spermatogonial differentiation. Antagonizing the activity of these two receptors with inhibitors, or a quadruple knockout of all adenosine receptors, increased the number of undifferentiated spermatogonia in both spermatogonial culture and *in vivo* germ cell development. Notably, this promoting effect on spermatogonial proliferation was abolished by inhibiting MEK activity, indicating a coordination between the adenosine and MEK-ERK pathways. In addition, adenosine may also affect other signaling pathways. For example, we observed that the addition of H89, a selective inhibitor of cAMP-dependent PKA ^27^, during differentiation, reduced the formation of KIT+ differentiating spermatogonia (Supplemental Fig. S4B). Transcript levels of differentiation markers *Stra8* and *Kit* were also downregulated upon H89 treatment (Supplemental Fig. S4C), suggesting that the activation of the PKA pathway favors spermatogonial differentiation. Adenosine and PKA pathways are known to be interconnected *via* cAMP ^13, 14^. Upregulated adenosine may increase cellular cAMP levels, thereby activating the PKA pathway to promote spermatogonial differentiation. Collectively, these findings indicate that the adenosine, MEK-ERK, and PKA pathways are intertwined in regulating spermatogonial fate decisions.

Our data suggest that adenosine functions in both the first wave and steady-state spermatogenesis, as we consistently observed smaller testes in *Ahcy* conditional knockout mice throughout germ cell development from prepuberty to adulthood. However, the fertility of *Ahcy* mutant mice appeared normal, likely because the transient impairment of spermatogenesis was not captured by the long-term fertility test. In addition, AHCY deletion cannot completely block adenosine synthesis, as adenosine can be generated from alternative pathways such as AMP dephosphorylation. Adenosine production from neighboring testicular interstitial cells may also activate adenosine receptors on remaining spermatogonia, promoting their survival and differentiation, thereby contributing to the recovery of spermatogenesis over time. Notably, quad-knockout mice with deletion of all 4 adenosine receptor genes exhibited a milder phenotype compared to *Ahcy* null mice. Transient spermatogenic arrest rapidly resolved around postnatal week 6.5 in QKO testes. It is possible that adenosine exerts multifunctional mechanisms to regulate spermatogonial development, only partially relying on acceptor-mediated pathways. Nevertheless, the balance between spermatogonial proliferation and differentiation is particularly sensitive to the changes in adenosine levels or receptor activities. We observed an increase in PLZF+ undifferentiated spermatogonia and a block in spermatogonial differentiation in mice either with *Ahcy* deletion or quad-adenosine receptor knockout. Together, our study has identified adenosine-mediated metabolic signaling as a previously underappreciated regulator that refines the balance between spermatogonial proliferation and differentiation. Given the conservation between humans and mice in germ cell development, our findings have broad implications for understanding the sensitivity of male germline stem cells to metabolic and environmental perturbations, thereby informing strategies to target metabolic pathways to improve reproductive health.

## Experimental Procedures

### Resource Availability

Lead Contact: Request for further information and resources should be directed to and fulfilled by Yuan Wang

Materials Availability: Requests for most unique reagents generated in this study should be directed to the lead contact to initiate a materials transfer agreement. Request for QKO mice of adenosine receptors should be addressed to Dr. Marc Reitman.

Data Availability: Metabolic data are provided in Supplemental Table S1.

### Mouse models used in this study

All animal experimental procedures were performed according to the approved protocol by the Institutional Animal Care and Use Committee at Michigan State University (PROTO202300165), NIH/NIDDK (K016-DEOB-23), and East China Normal University (m20200902). *Ahcy* conditional mice were purchased from GemPharmatech Co. Ltd., China. Founder mice with the correct floxed alleles were identified by PCR-based genotyping and then crossed with C57BL/6 mice (000664; the Jackson Laboratory) to generate heterozygous *Ahcy^+/f^* mice. *Ahcy^f/f^; Cre* were obtained by crossing heterozygote *Ahcy^+/f^; Cre* males with *Ahcy^f/f^* female mice. Adenosine receptor QKO ^12^ and *Stra8-Cre* ^17^ mice were previously described.

### Histology Study, IHF, and IF Assays

Testes were fixed in Bouin’s solution (Sigma-Aldrich, HT10132) or 4% paraformaldehyde, followed by dehydration, embedding in paraffin, and cut into 4 µm sections for hematoxylin (Sigma-Aldrich, MHS1) and eosin (Sigma-Aldrich, E4009) staining. For histology, permount (SP15100; Fisher Scientific) was applied onto the sections. The sections were then covered by coverslips and dried in the hood overnight before imaging. For IHF staining, 4 µm testis sections were permeabilized with PBS containing 0.1% Triton X-100 and washed with PBS. 10 mM sodium citrate buffer (pH = 6.0) was used for antigen retrieval. Sections were blocked with in PBS containing 5% BSA for 1 hour at room temperature, followed by incubation with primary antibodies overnight at 4°C. After washing with PBS, sections were incubated with fluorophore-conjugated secondary antibodies for 1 hour at room temperature in the dark. Sections were mounted with Glass Antifade Mountant (P36982; Invitrogen) for imaging. Antibodies were used in this study below: PRM1 (Briar Patch Biosciences, MAb-Hup1N-150, 1:200), Antibodies from SANTA CRUZ: PLZF (SC-28319,1:500), AHCY (SC-271389,1:500), Antibodies from Abcam: DDX4 (ab13840 1:500), SYCP3 (ab15093,1:500), STRA8 (Ab49602 and 49405,1:500). Antibodies from Invitrogen: Donkey anti-Mouse Alexa Fluor 488 (A21202,1:500), Donkey anti-Rabbit Alexa Fluor 488 (A32790TR,1:500), Donkey anti-Rabbit Alexa Fluor 647 (A-31573,1:500). Antibodies from Jackson ImmunoResearch: Goat anti-Mouse Alexa Fluor 594 (115-585-003,1:500), Goat anti-Rabbit Alexa Fluor 594 (111-585-003,1:500). Microscopes were used in this study: Olympus FL confocal microscope, Nikon Eclipse Ti microscope and BioTek Lionheart FX Automated Microscope, Leica DM4000B microscope, and Leica TCS/SP5 confocal microscope.

### Computer-Assisted Sperm Analysis

The cauda epididymis was dissected in PBS. The tissue was gently minced with micro-scissors to release sperm and incubated in HTF medium at 37°C for about 15 min. Sperm concentration was determined using a computer-assisted sperm analysis system (Hamilton Thorne IVOS II), following the Manufacturer’s instructions.

### Cell Culture, Transfection, Viral Infection, and RNA interference by shRNAs

Mouse spermatogonial culture was established from postnatal day 5-7 testes of DBA/2J mice or from the testes of F1 progeny derived from a CAG-EGFP line (The Jackson Laboratory, 003291) crossed with DBA/2J mice. Undifferentiated spermatogonia were isolated using a FITC-tagged CD9 antibody (124808; BioLegend) with a Miltenyi Biotec MACS separation kit, according to the manufacturer’s instructions, and cultured in SSC culture medium, as described in published protocols ^28, 29^. For packaging virus with shRNA expression, HEK293T cells were cultured in DMEM (Gibco) with 10% fetal bovine serum (Gibco) and transfected with pll3.7-U6 (Addgene plasmid # 11795) expressing target-specific shRNAs and psPAX2 and pMD2.G packaging plasmids into HEK293T cells using polyethyleneimine (Sigma). For viral infection of spermatogonia, lentivirus were concentrated by Amicon® Ultra-50 Centrifugal Filter Unit (Sigma, UFC9100) into 6X10^7^/ml. Cell suspension was mixed with concentrated virus with MOI at 2 and 5 µg/mL polybrene. Cell medium was changed after 20 hours post infection. Gene knockdown efficiency was verified by real-time RT-PCR. Small molecules used in culture: adenosine (Sigma, 100 nM); ZM241385 (SelleckChem, 2 µM); GS6201 (MedChemExpress, 2 µM); PD0325091 (SelleckChem, 1 µM). Oligos for shRNA knockdown are listed in Supplemental Table S2.

### HILIC-LC-MS/MS Analysis

For sample preparation, 1 × 10⁶ spermatogonia per well were plated onto gelatin-coated 6-well plates. After 24 hours, cells were treated with either 10 nM RA or an equal volume of DMSO as controls. Cells were harvested for targeted metabolomic analysis after 48-hour treatment. Six biological replicates were used and normalized to the total protein levels. Targeted metabolomic analysis was performed using hydrophilic interaction liquid chromatography coupled with tandem mass spectrometry (HILIC-LC-MS/MS). Briefly, metabolites were extracted from cells using methanol/acetonitrile/water (2:2:1, v/v), with ultrasonication, followed by protein precipitation at −20°C for 1 hour and centrifugation at 14,000g, 4°C. The supernatants were lyophilized and reconstituted in acetonitrile/water (1:1, v/v) for analysis. Chromatographic separation was achieved on a Waters ACQUITY UPLC BEH Amide column (2.1 × 100 mm, 1.7 μm) with a binary gradient of mobile phase A (25 mM ammonium acetate and 25 mM ammonium hydroxide, pH ∼9.75) and mobile phase B (acetonitrile). Metabolites were quantified using multiple reaction monitoring (MRM) on an AB Sciex 5500 QTRAP mass spectrometer, with MRM parameters optimized (sheath gas temperature, 350°C; dry gas temperature, 350°C; sheath gas flow, 11L/min; dry gas flow, 10 L/min; capillary voltage, 4000 V in positive mode and -3500 V in negative modes respectively; nozzle voltage, 500 V; and nebulizer pressure, 30 psi). Quantification was performed using authentic standards for metabolite identification and peak area integration. Quality control was maintained through pooled QC samples, accepting metabolites with RSD < 30%. Data were processed with MRMAnalyzer, and differential metabolite levels in undifferentiated and differentiating spermatogonia were determined by Student’s t-test with *p* < 0.05 and fold change > 1.5. Raw data were uploaded to Metabolomics Workbench Repository at: https://dev.metabolomicsworkbench.org:22222/data/DRCCMetadata.php?Mode=Study&StudyID=ST004924&Access=ZqhR9558

### Flow Cytometry

Testes or cultured cells were digested to a single-cell suspension and stained in the dark with appropriate dyes or antibodies: BV421-KIT (561074; BD Biosciences) and PE-CD90 (105308; BioLegend). Flow cytometers used in this study were: BD LSRII, Cytek Aurora, and Attune CytPix.

### Western Blot

Western blots were performed according to standard protocols. Primary antibodies were used in this study: p-ERK1/2 (Cell Signaling Technology, 4370S, 1:200), ERK1/2 (Cell Signaling Technology, 4696S, 1:500), β-Actin (Sigma, A5441,1:5000). The fluorochrome-conjugated 2^nd^ antibodies (LI-COR Biosciences, P/N: 926-49020; LI-COR Biosciences P/N: 926-68070) were used at 1:10000 dilution. The fluorescent signals against targeted proteins were detected using the Li-COR Odyssey system.

### Statistical Analysis

Data were presented as mean ± SEM. All experiments were performed independently three times or more unless otherwise stated. Statistical comparisons between-group means were performed using the unpaired Student’s t-test or one-way ANOVA followed by Tukey’s post-hoc test in the GraphPad Prism software.

## Acknowledgements

This work was partially supported by the grants from the National Institutes of Health and the National Institute of General Medical Sciences (GM146587) and Hatch Fund (MICL02781) from the US Department of Agriculture (USDA) and the National Institute of Food and Agriculture (NIFA) *via* MSU AgbioResearch to Dr. Yuan Wang. The Attune CytPix at the MSU flow cytometry used in this study is supported by the Equipment Grants Program (award #: 2022-70410-38419) from the USDA and NIFA. We thank Dr. Minghan Tong at the Chinese Academy of Sciences for providing *Stra8-Cre* mice. This research was supported in part by the Intramural Research Program of the National Institute of Diabetes and Digestive and Kidney Diseases (NIDDK) within the National Institutes of Health (NIH). The contributions of the NIH author(s) are considered Works of the United States Government. The findings and conclusions presented in this paper are those of the author(s) and do not necessarily reflect the views of the NIH or the U.S. Department of Health and Human Services.

## Author Contributions

Wang P, Nguyen D, Xu Y, Hu B, and Zhang Z performed the experiments and analyzed data. Reitman M and Xiao C provided QKO testes and edited manuscript. Wang Y designed experiments and analyzed data. Wang P and Wang Y wrote the manuscript.

## Author Contributions

Wang P, Nguyen D, Xiao C, Xu Y, Hu B, and Zhang Z performed the experiments and analyzed data. Reitman M provided resources. Wang P and Wang Y designed experiments, analyzed data, and wrote the manuscript.

## Supplemental Figure Legends

**Fig. S1.**
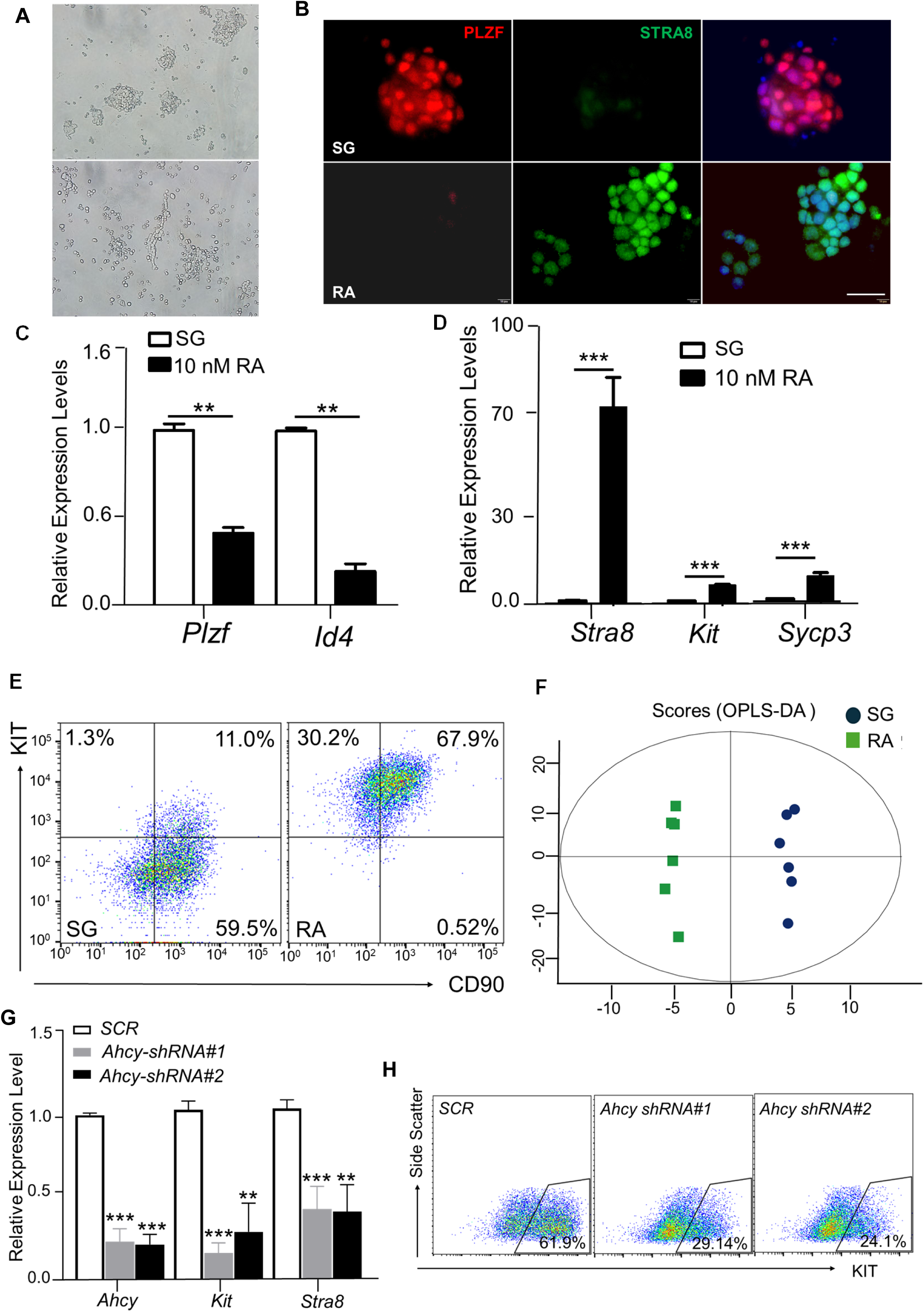
Adenosine level is increased during spermatogonial differentiation. (**A-B**) Images of cultured spermatogonia with RA (RA) or without RA (SG) induction under a bright-field microscope (**A**) and were stained with antibodies against the undifferentiated marker PLZF and the differentiating marker STRA8, counterstained with the nuclear dye DAPI (**B**). (**C-D**) The markers for undifferentiated (PLZF and ID4) and differentiating (*Stra8, Kit,* and *Sycp3*) spermatogonia were analyzed by real-time RT-PCR. Data were shown as mean ± SEM. **: *p* <0.01; ***: *p* < 0.001; n=4. (**E**) Cultured spermatogonia were induced to differentiate with RA. The undifferentiated spermatogonia (SG) and RA-induced differentiating population were analyzed by flow cytometry using antibodies against CD90 and KIT. (**F**) Orthogonal Projections to Latent Structures – Discriminant Analysis of Metabolome Data. 6 biological replicates per group were included and showed well-separated clustering of SG and RA-treated samples. (**G**) The expression levels of differentiation markers were analyzed by real-time RT-PCR in RA-induced differentiating spermatogonia treated with scrambled shRNA (SCR) or two different *Ahcy* shRNAs. *Ahcy* knockdown led to decreased *Kit* and *Stra8* expression upon RA induction compared with the SCR control. Data were shown as mean ± SEM. N=3. **: *p* <0.01; ***: *p* < 0.001. (**H**) Spermatogonia upon *Ahcy* knockdown were induced by RA and analyzed by flow cytometry. A decreased percentage of KIT+ differentiating spermatogonia was observed.

**Fig. S2.**
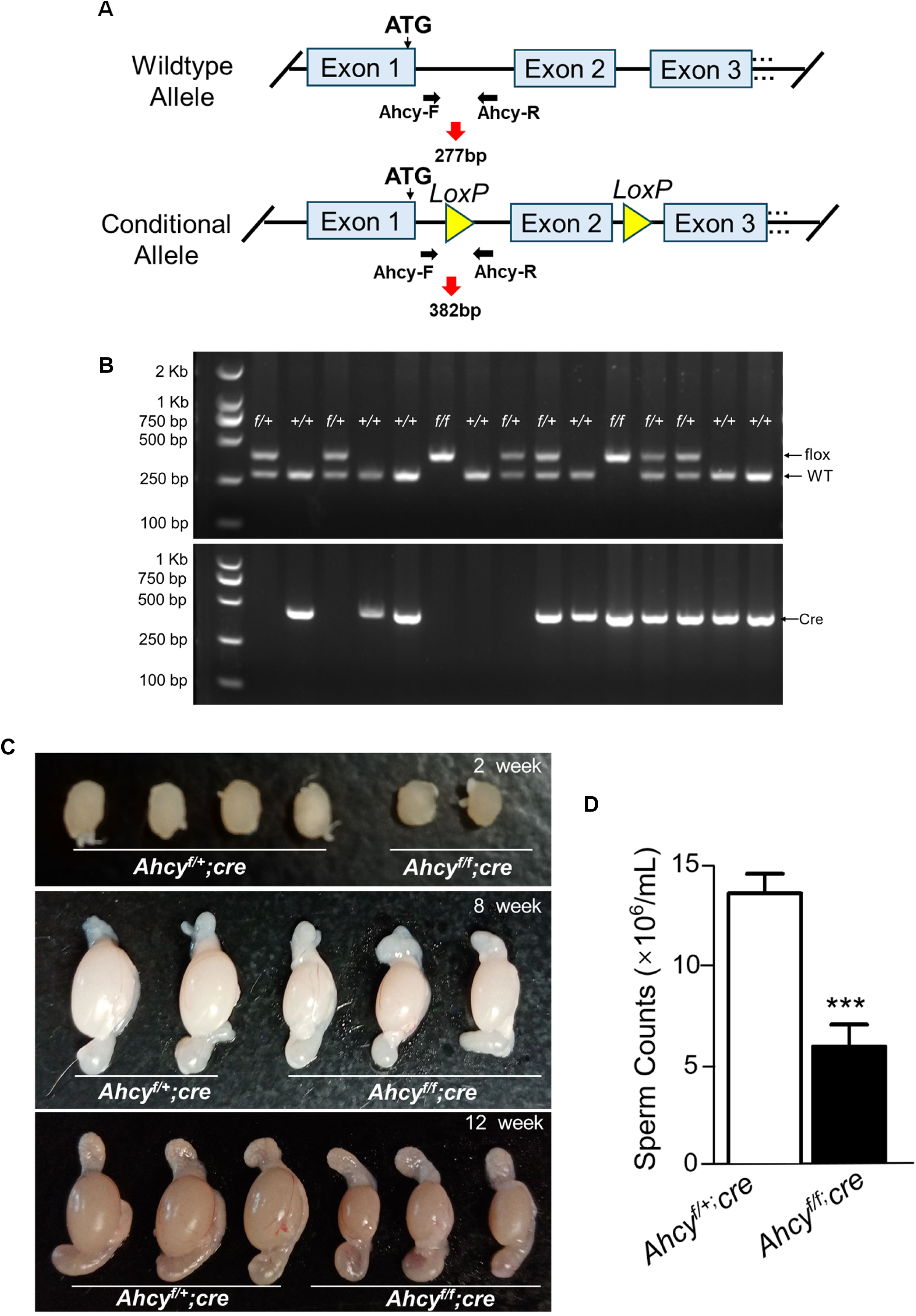
Establishing an *Ahcy* conditional knockout mouse model. **(A)** A diagram shows the *Ahcy* knockout strategy. Exon 2 was flanked by two *LoxP* sites within the modified *Ahcy* allele, leading to out-of-frame *Ahcy* coding sequences. **(B)** An example illustrates the genotyping results using PCR analysis on *Ahcy^f/f^* and wild-type mouse alleles. The PCR with the primer pair Ahcy-F and Ahcy-R amplifies the wild-type *Ahcy* allele with a 277 bp product and the floxed *Ahcy* allele at 382 bp. The band in *the Stra8-Cre* PCR shows the existence of the Cre transgene. **(C)** The gross morphology of testes and epididymis from *Ahcy^f/f^; Cre* mice and control littermates at various ages. **(D)** The sperm concentration from epididymis of mice at 8 week old was measured by CASA. Data were shown as mean ± SEM. ***: *p* < 0.001. n=5.

**Fig. S3.**
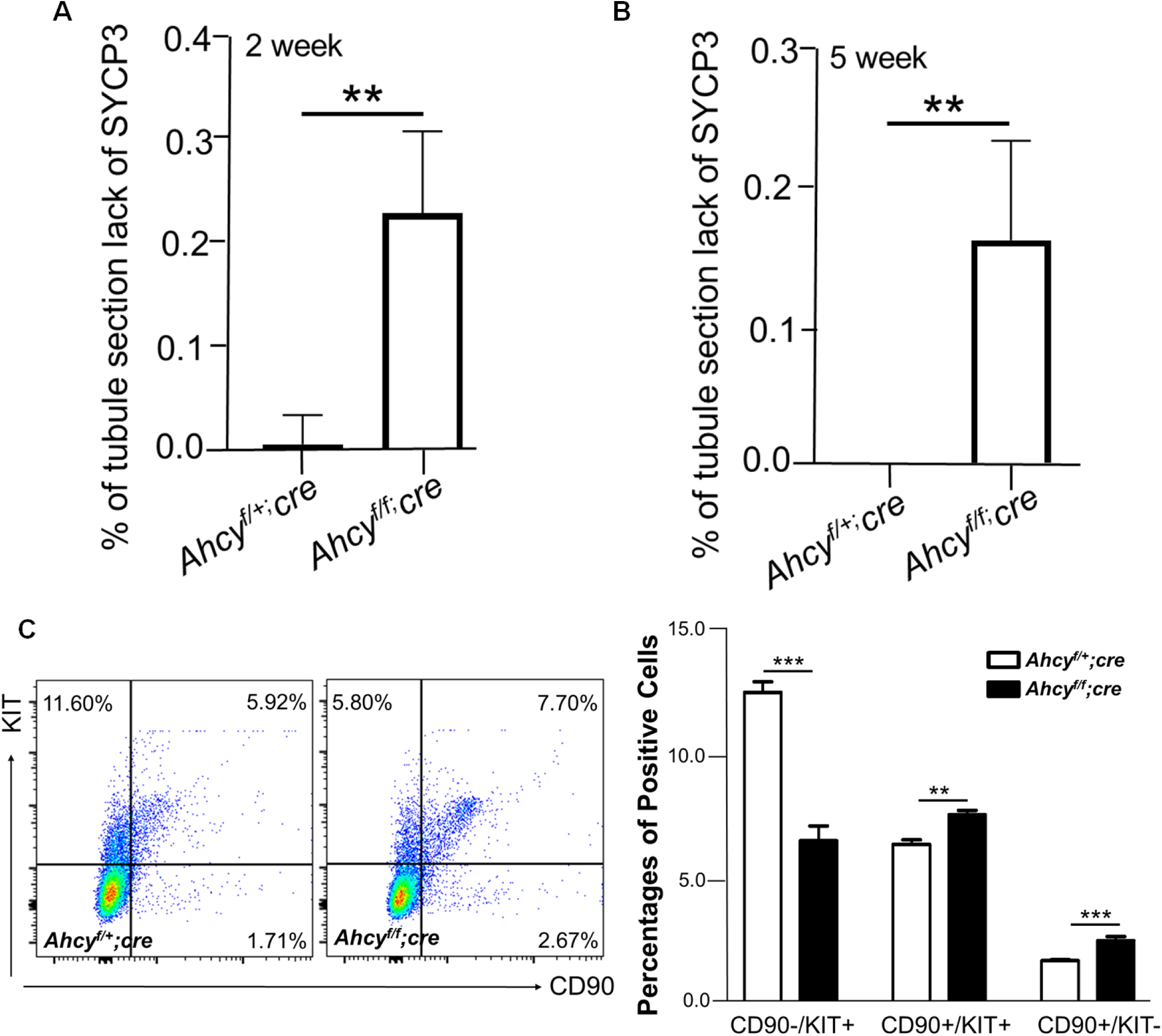
*Ahcy* conditional knockout delays spermatogenesis. **(A-B)** The percentages of seminiferous tubule section lack of SYCP3+ spermatocytes, calculated from IHF on *Ahcy^f/f^; Cre* (93 tubule sections from 13 slides) and littermate control (112 tubule sections from 20 slides) testes from 2-week-old (**A**) and 5-week-old (**B**) mice. (**C**) Flow cytometry analyses on *Ahcy^f/f^; Cre* and control testes. Left Panel: representative images of flow cytometry. Right Panel: A bar graph shows the average percentages of undifferentiated (CD90+/KIT-) and CD90-/KIT+ differentiating spermatogonia in *Ahcy^f/f^; Cre* and control testes, calculated from 3 littermate paired mice. (**A-C**) Data were shown as mean ± SEM. **: *p* <0.01. ; ***: *p* < 0.001.

**Fig. S4.**
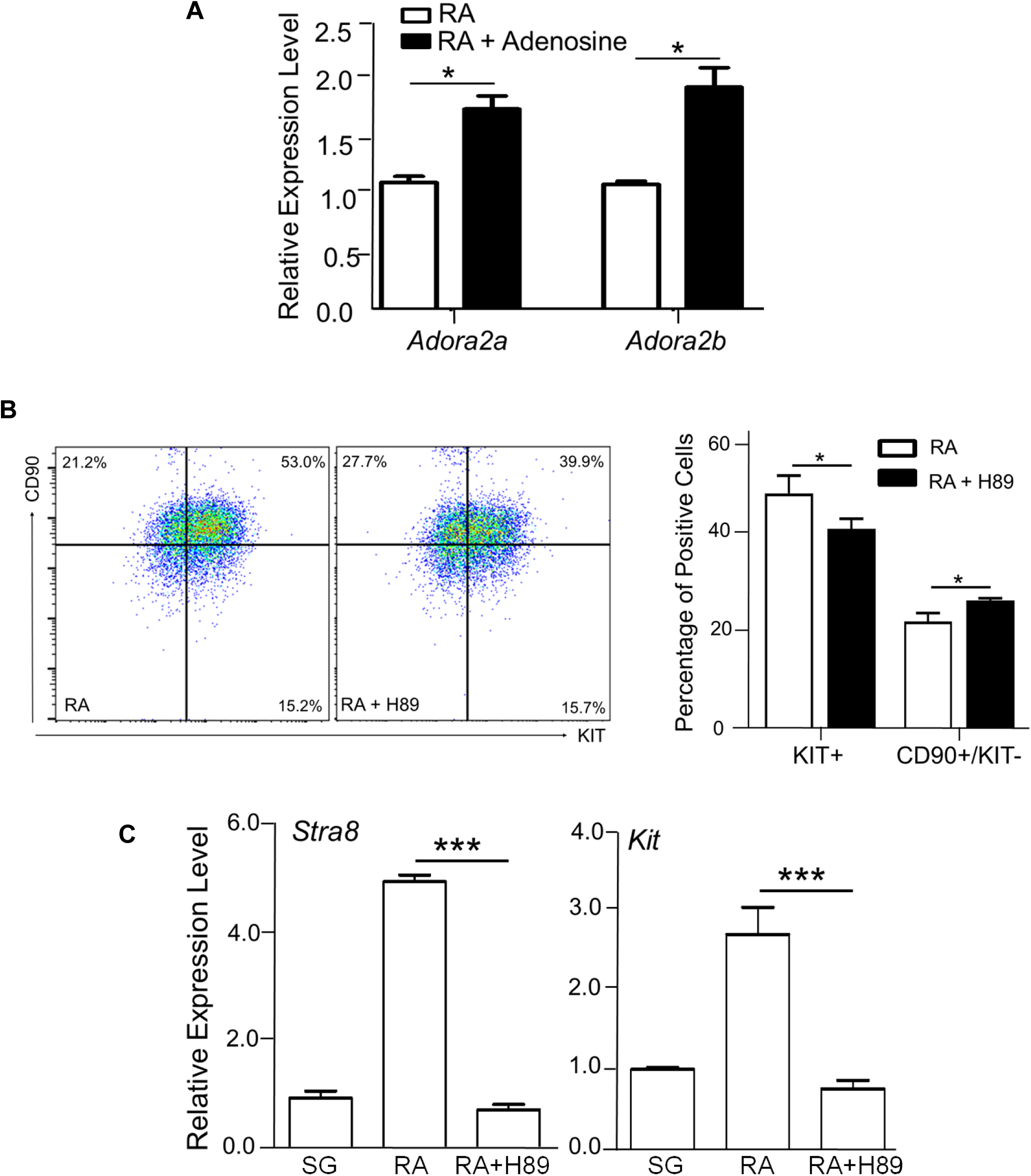
Adenosine receptor activities in spermatogonial development. (**A**) Spermatogonia were induced to differentiate by RA in the absence or presence of adenosine treatment. The transcript levels of adenosine receptor *A2a* and *A2b* were analyzed using real-time RT-PCR. (**B**) Flow cytometry analyses of CD90 and KIT on RA-induced spermatogonia in the absence or presence of a PKA inhibitor, H89. Left panel: representative flow cytometry graphs. Right panel: a bar graph shows the average percentages of CD90+/KIT-undifferentiated and KIT+ differentiated spermatogonia. (**C**) Expression levels of differentiation markers were analyzed by real-time RT-PCR in undifferentiated (SG) and differentiating spermatogonia treated with RA or RA plus H89. **(A-C)** Data were shown as mean ± SEM. *: *p* <0.05; **: *p* < 0.01; ***: *p* < 0.001. N=3.

**Fig. S5.** Raw unprocessed Western Blot image to support Fig. 4D.

**Fig. S6.** Raw unprocessed gel image to support Fig. S2B.

## Supplemental Tables

**Supplemental Table S1.** Levels of metabolites detected in undifferentiated spermatogonia and differentiating spermatogonia induced by RA.

**Supplemental Table S2.** Sequences of oligonucleotides used in this study.

## References

1. Griswold MD, Oatley JM. Concise review: Defining characteristics of mammalian spermatogenic stem cells. Stem Cells. 2013;31(1):8–11. doi: 10.1002/stem.1253. PubMed PMID: 23074087; PMCID: PMC5312674.

2. Mecklenburg JM, Hermann BP. Mechanisms Regulating Spermatogonial Differentiation. Results Probl Cell Differ. 2016;58:253–87. Epub 2016/06/15. doi: 10.1007/978-3-319-31973-5_10. PubMed PMID: 27300182.

3. Nakamura M, Fujiwara A, Yasumasu I, Okinaga S, Arai K. Changes in Activities of Glucose Metabolising Enzymes in Germ Cells during Spermatogenesis in Rats: (glucose-6-phosphate dehydrogenase/germ cells/spermatogenesis/rat). Dev Growth Differ. 1983;25(3):281–7. Epub 1983/01/01. doi: 10.1111/j.1440-169X.1983.00281.x. PubMed PMID: 37282215.

4. Nakamura M, Okinaga S, Arai K. Metabolism of pachytene primary spermatocytes from rat testes: pyruvate maintenance of adenosine triphosphate level. Biol Reprod. 1984;30(5):1187–97. Epub 1984/06/01. doi: 10.1095/biolreprod30.5.1187. PubMed PMID: 6733209.

5. Chen W, Zhang Z, Chang C, Yang Z, Wang P, Fu H, Wei X, Chen E, Tan S, Huang W, Sun L, Ni T, Yang Y, Wang Y. A bioenergetic shift is required for spermatogonial differentiation. Cell Discov. 2020;6:56. Epub 2020/08/31. doi: 10.1038/s41421-020-0183-x. PubMed PMID: 32864161; PMCID: PMC7431567.

6. Kanatsu-Shinohara M, Tanaka T, Ogonuki N, Ogura A, Morimoto H, Cheng PF, Eisenman RN, Trumpp A, Shinohara T. Myc/Mycn-mediated glycolysis enhances mouse spermatogonial stem cell self-renewal. Genes Dev. 2016;30(23):2637–48. Epub 2016/12/23. doi: 10.1101/gad.287045.116. PubMed PMID: 28007786; PMCID: PMC5204355.

7. Lord T, Law NC, Oatley MJ, Miao D, Du G, Oatley JM. A novel high throughput screen to identify candidate molecular networks that regulate spermatogenic stem cell functionsdagger. Biol Reprod. 2022;106(6):1175–90. Epub 2022/03/05. doi: 10.1093/biolre/ioac048. PubMed PMID: 35244684; PMCID: PMC9198950.

8. Varuzhanyan G, Rojansky R, Sweredoski MJ, Graham RLJ, Hess S, Ladinsky MS, Chan DC. Mitochondrial fusion is required for spermatogonial differentiation and meiosis. Elife. 2019;8. Epub 2019/10/10. doi: 10.7554/eLife.51601. PubMed PMID: 31596236; PMCID: PMC6805159.

9. Boison D, Yegutkin GG. Adenosine Metabolism: Emerging Concepts for Cancer Therapy. Cancer Cell. 2019;36(6):582–96. Epub 2019/12/11. doi: 10.1016/j.ccell.2019.10.007. PubMed PMID: 31821783; PMCID: PMC7224341.

10. Zhulai G, Oleinik E, Shibaev M, Ignatev K. Adenosine-Metabolizing Enzymes, Adenosine Kinase and Adenosine Deaminase, in Cancer. Biomolecules. 2022;12(3). Epub 2022/03/26. doi: 10.3390/biom12030418. PubMed PMID: 35327609; PMCID: PMC8946555.

11. Chen JF, Eltzschig HK, Fredholm BB. Adenosine receptors as drug targets--what are the challenges? Nat Rev Drug Discov. 2013;12(4):265–86. Epub 2013/03/29. doi: 10.1038/nrd3955. PubMed PMID: 23535933; PMCID: PMC3930074.

12. Xiao C, Liu N, Jacobson KA, Gavrilova O, Reitman ML. Physiology and effects of nucleosides in mice lacking all four adenosine receptors. PLoS Biol. 2019;17(3):e3000161. Epub 2019/03/02. doi: 10.1371/journal.pbio.3000161. PubMed PMID: 30822301; PMCID: PMC6415873.

13. Vincenzi F, Pasquini S, Contri C, Cappello M, Nigro M, Travagli A, Merighi S, Gessi S, Borea PA, Varani K. Pharmacology of Adenosine Receptors: Recent Advancements. Biomolecules. 2023;13(9). Epub 2023/09/28. doi: 10.3390/biom13091387. PubMed PMID: 37759787; PMCID: PMC10527030.

14. Raskovalova T, Lokshin A, Huang X, Jackson EK, Gorelik E. Adenosine-mediated inhibition of cytotoxic activity and cytokine production by IL-2/NKp46-activated NK cells: involvement of protein kinase A isozyme I (PKA I). Immunol Res. 2006;36(1-3):91–9. Epub 2007/03/06. doi: 10.1385/IR:36:1:91. PubMed PMID: 17337770.

15. Migita H, Kominami K, Higashida M, Maruyama R, Tuchida N, McDonald F, Shimada F, Sakurada K. Activation of adenosine A1 receptor-induced neural stem cell proliferation via MEK/ERK and Akt signaling pathways. J Neurosci Res. 2008;86(13):2820–8. Epub 2008/07/12. doi: 10.1002/jnr.21742. PubMed PMID: 18618669.

16. Vaccaro JA, Naser SA. The Role of Methyl Donors of the Methionine Cycle in Gastrointestinal Infection and Inflammation. Healthcare (Basel). 2021;10(1). Epub 2022/01/22. doi: 10.3390/healthcare10010061. PubMed PMID: 35052225; PMCID: PMC8775811.

17. Lin Z, Hsu PJ, Xing X, Fang J, Lu Z, Zou Q, Zhang KJ, Zhang X, Zhou Y, Zhang T, Zhang Y, Song W, Jia G, Yang X, He C, Tong MH. Mettl3-/Mettl14-mediated mRNA N(6)-methyladenosine modulates murine spermatogenesis. Cell Res. 2017;27(10):1216–30. Epub 2017/09/16. doi: 10.1038/cr.2017.117. PubMed PMID: 28914256; PMCID: PMC5630681.

18. Jaakola VP, Griffith MT, Hanson MA, Cherezov V, Chien EY, Lane JR, Ijzerman AP, Stevens RC. The 2.6 angstrom crystal structure of a human A2A adenosine receptor bound to an antagonist. Science. 2008;322(5905):1211–7. Epub 2008/10/04. doi: 10.1126/science.1164772. PubMed PMID: 18832607; PMCID: PMC2586971.

19. Xu F, Wu H, Katritch V, Han GW, Jacobson KA, Gao ZG, Cherezov V, Stevens RC. Structure of an agonist-bound human A2A adenosine receptor. Science. 2011;332(6027):322–7. Epub 2011/03/12. doi: 10.1126/science.1202793. PubMed PMID: 21393508; PMCID: PMC3086811.

20. Toldo S, Zhong H, Mezzaroma E, Van Tassell BW, Kannan H, Zeng D, Belardinelli L, Voelkel NF, Abbate A. GS-6201, a selective blocker of the A2B adenosine receptor, attenuates cardiac remodeling after acute myocardial infarction in the mouse. J Pharmacol Exp Ther. 2012;343(3):587–95. Epub 2012/08/28. doi: 10.1124/jpet.111.191288. PubMed PMID: 22923737; PMCID: PMC11047795.

21. Hasegawa K, Namekawa SH, Saga Y. MEK/ERK signaling directly and indirectly contributes to the cyclical self-renewal of spermatogonial stem cells. Stem Cells. 2013;31(11):2517–27. Epub 2013/07/31. doi: 10.1002/stem.1486. PubMed PMID: 23897718; PMCID: PMC3834200.

22. He Z, Jiang J, Kokkinaki M, Golestaneh N, Hofmann MC, Dym M. Gdnf upregulates c-Fos transcription via the Ras/Erk1/2 pathway to promote mouse spermatogonial stem cell proliferation. Stem Cells. 2008;26(1):266–78. Epub 2007/10/27. doi: 10.1634/stemcells.2007-0436. PubMed PMID: 17962702; PMCID: PMC2905627.

23. Schulte G, Fredholm BB. Human adenosine A(1), A(2A), A(2B), and A(3) receptors expressed in Chinese hamster ovary cells all mediate the phosphorylation of extracellular-regulated kinase 1/2. Mol Pharmacol. 2000;58(3):477–82. Epub 2000/08/23. PubMed PMID: 10953039.

24. Dura M, Teissandier A, Armand M, Barau J, Lapoujade C, Fouchet P, Bonneville L, Schulz M, Weber M, Baudrin LG, Lameiras S, Bourc’his D. DNMT3A-dependent DNA methylation is required for spermatogonial stem cells to commit to spermatogenesis. Nat Genet. 2022;54(4):469–80. Epub 2022/04/13. doi: 10.1038/s41588-022-01040-z. PubMed PMID: 35410378.

25. Xiao Z, Locasale JW. Epigenomic links from metabolism-methionine and chromatin architecture. Curr Opin Chem Biol. 2021;63:11–8. Epub 2021/03/06. doi: 10.1016/j.cbpa.2021.01.011. PubMed PMID: 33667809; PMCID: PMC9889272.

26. Zhou J, Yang L, Zhong T, Mueller M, Men Y, Zhang N, Xie J, Giang K, Chung H, Sun X, Lu L, Carmichael GG, Taylor HS, Huang Y. H19 lncRNA alters DNA methylation genome wide by regulating S-adenosylhomocysteine hydrolase. Nat Commun. 2015;6:10221. Epub 2015/12/22. doi: 10.1038/ncomms10221. PubMed PMID: 26687445; PMCID: PMC4703905.

27. Lochner A, Moolman JA. The many faces of H89: a review. Cardiovasc Drug Rev. 2006;24(3-4):261–74. Epub 2007/01/12. doi: 10.1111/j.1527-3466.2006.00261.x. PubMed PMID: 17214602.

28. Kanatsu-Shinohara M, Ogonuki N, Inoue K, Miki H, Ogura A, Toyokuni S, Shinohara T. Long-term proliferation in culture and germline transmission of mouse male germline stem cells. Biol Reprod. 2003;69(2):612–6. Epub 2003/04/18. doi: 10.1095/biolreprod.103.017012. PubMed PMID: 12700182.

29. Zhang Z, Miao J, Wang H, Ali I, Nguyen D, Chen W, Wang Y. Accelerated mitochondrial dynamics promote spermatogonial differentiation. Stem Cell Reports. 2024;19(11):1548–63. Epub 2024/10/12. doi: 10.1016/j.stemcr.2024.09.006. PubMed PMID: 39393359; PMCID: PMC11589200.

